# Genotypic context modulates fitness landscapes: Effects on the speed and direction of evolution for antimicrobial resistance

**DOI:** 10.1101/427328

**Authors:** C. Brandon Ogbunugafor, Rafael F. Guerrero, Margaret J. Eppstein

## Abstract

Understanding the forces that drive the dynamics of adaptive evolution is a goal of many subfields within evolutionary biology. The fitness landscape analogy has served as a useful abstraction for addressing these topics across many systems, and recent treatments have revealed how different environments can frame the particulars of adaptive evolution by changing the topography of fitness landscapes. In this study, we examine how the larger, ambient genotypic context in which the fitness landscape being modeled is embedded affects fitness landscape topography and subsequent evolution. Using simulations on empirical fitness landscapes, we discover that genotypic context, defined by genetic variability in regions outside of the locus under study (in this case, an essential bacterial enzyme target of antibiotics), influences the speed and direction of evolution in several surprising ways. These findings have implications for how we study the evolution of drug resistance in nature, and for presumptions about how biological evolution might be expected to occur in genetically-modified organisms. More generally, the findings speak to theory surrounding how “difference can beget difference” in adaptive evolution: that small genetic differences between organisms can greatly alter the specifics of how evolution occurs, which can rapidly drive even slightly diverged populations further apart.

**Author summary:** Technological advances enable scientists to engineer individual mutations at specific sites within an organism’s genome with increasing ease. These breakthroughs have provided scientists with tools to study how different engineered mutations affect the function of a given gene or protein, yielding useful insight into genotype-phenotype mapping and evolution. In this study, we use engineered strains of bacteria to show how the dynamics (speed and direction) of evolution of drug resistance in an enzyme depends on the species-type of that bacterial enzyme, and on the presence/absence of mutations in other genes in the bacterial genome. These findings have broad implications for public health, genetic engineering, and theories of speciation. In the context of public health and biomedicine, our results suggest that future efforts in managing antimicrobial resistance must consider genetic makeup of different pathogen populations before predicting how resistance will occur, rather than assuming that the same resistance pathways will appear in different pathogen populations. With regard to broader theory in evolutionary biology, our results show how even small genetic differences between organisms can alter how future evolution occurs, potentially causing closely-related populations to quickly diverge.

## Introduction

The fitness landscape analogy has undergone a subtle makeover in recent years, with larger data sets and improved methods (laboratory and computational) greatly increasing the scope of systems and questions that the analogy can be used to responsibly address. For example, recent studies have examined how environments change adaptive landscape topography [1, 2], employed methods to construct adaptive landscapes in natural populations [3, 4], and conducted large scale examinations of epistasis acting across fitness landscapes [5–10]. Other examinations have extracted new information out of empirical fitness landscapes, including how landscapes changes in shape during adaptive evolution [11], how indirect pathways are traversed during evolution [12], and how features of a landscape determine the speed of some adaptive trajectories relative to others [13]. The theme across many of these recent breakthroughs is a growth in our understanding of how various contexts can frame our expectations for how evolution will occur, and render it challenging to predict [14–17]. This is of particular importance in studies utilizing empirically determined fitness landscapes to understand the evolution of drug resistance, where the hope is to one day understand how the evolution of resistance occurs such that disease can be treated more effectively [18–20]

Importantly, specific portions of the genome that are the object of study in model systems (e.g., a single gene encoding a single protein) do not function in genetic isolation in natural settings. Rather, findings affirming that mutations and loci throughout the genome often interact (sometimes in a non-linear fashion) can now be considered the norm in modern evolutionary genetics. Thus, it is very likely that portions of the genome that are not the object of study (i.e., those defining the broader “genotypic context” of the organism) can alter the portion of the empirical fitness landscape under study. And insofar as genotypic context complicates genotype-phenotype mapping in general, it may also play a role in crafting how adaptive evolution occurs. We can quickly recapitulate this intrigue with a simple question: if we engineer the same mutation into two different strains of an organism, how would we expect the genetic differences between these organisms to influence downstream evolution to a common stressor in each strain?

In this study, we directly examine empirical fitness landscapes constructed for the study of antimicrobial resistance in dihydrofolate folate reductase (DHFR), an essential bacterial enzyme. Specifically, we employ a data set whereby three orthologous mutations associated with drug resistance were engineered (in all eight possible combinations) into DHFRs from three species of bacteria (*Escherichia coli, Listeria grayi*, and *Chlamydia muridarum*). In addition, each of these alleles were then engineered into background bacterial strains containing three different protein quality control (PQC) profiles: wild type, GroEL chaperonins overexpression (GroEL+), and Lon protease knockout (Δ*lon*) [21]. This amounts to nine different genotypic contexts for the eight possible genotypes for these three DHFR loci associated with drug resistance. *IC*_50_ values were measured for the drug Trimethoprim (an anti-folate drug that targets DHFR) resulting in nine distinct empirical fitness landscapes. Using inferred growth rates based on the laboratory-derived *IC*_50_ values (see Materials and methods), we simulate evolution across all landscapes, identifying which paths are preferred in each landscape, measuring how many generations are required for the terminal genotype to become dominant (*T_d_*) and calculating the within-path competition (*C_w_*), a metric that has been shown to govern the speed of evolution across a trajectory [13] under standard assumptions of replicator dynamics [22]. Our findings are striking beyond the basic observation that genotypic context frames adaptive evolution. We show that the speed of adaptation differs drastically between contexts, and in a pattern that defies our intuition. For example, in modeling the evolution of resistance to Trimethoprim, we find that off-target mutations influence the landscape as much or more than those within the actual drug target. We discuss these findings with regard to how they speak to our efforts at modeling drug resistance, and more generally, how they affect our understanding of which forces craft how populations diverge.

## Materials and methods

### Empirical Fitness Landscapes

We utilized data that were previously generated to determine the biophysical components of a fitness landscape for resistance [21]. From it, we utilized *IC*_50_ values for the antibiotic Trimethoprim, for the eight genotypes with or without each of three mutant alleles (L28R, A26T, and P21L) in the DHFR gene, in three species of bacteria (*E. coli, L. grayi, C. muridarum*), each with three PQC profiles (wild type, GroEL+, or Δ*lon*) [21]. Although the number of replicate measurements for each genotype varied from two to six, for consistency we averaged only the first two replicates for each genotype, and report these values in Tables 1,2,3. We then inferred bacterial growth rates at high dosages of Trimethoprim from these *IC*_50_ values, as follows. We first verified that *IC*_50_ values are strongly correlated with growth rates at very high drug dosage by regressing published growth rates [2, 13] for 16 genotypes of *P. falciparum* DHFR exposed to high (10^5^ *μM*) dosages of Pyrimethamine and Cycloguanil against their published *IC*_50_ values [23, 24] and observed a very strong correlation (*R*^2^ > 0.99, *p* < 10^-35^). We then used the resulting regression equation to infer relative growth rates at 10^5^*μM* of Trimethoprim from the *IC*_50_ averages, as shown in Tables 1,2,3. Each of these nine sets of eight inferred growth rates (for the eight genotypes of a given species in a given PQC genetic profile) comprise a fitness landscape, as illustrated in Fig. 1. Note that four of these nine small fitness landscapes contain suboptimal peaks, reflecting the highly epistatic interactions between the three mutations [21].

**Fig 1.**
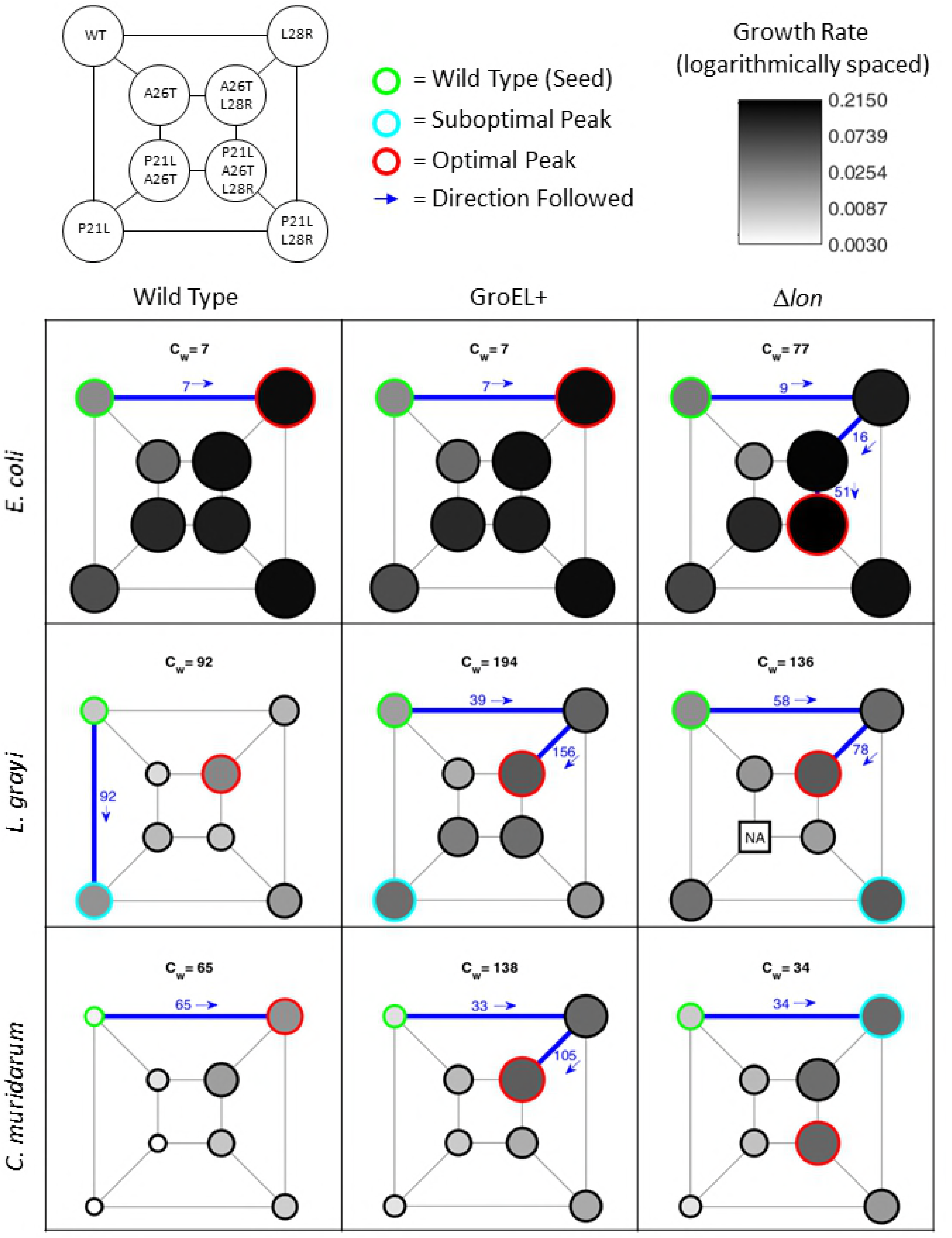
Fitness landscapes. Empirical fitness landscapes for the three species (rows) with three genetic backgrounds (columns) exposed to 10^5^*μM* of Trimethoprim. Nodes represent the genotypes indicated in the upper left diagram, where edges connect single-mutational neighbors. Node diameters and shading are proportional to the logarithm of the growth rates shown in Tables 1–3 (no growth rates were available for the square node labeled NA). Simulations (e.g., as shown in Fig. 2) starting from the wild type (WT, circled in green) follow the 1–3 step trajectories shown by the thick blue edges; each edge is labeled with the within-path competition (*C_w_*) for that step and the *C_w_* for the entire trajectory is shown above each landscape. Each trajectory terminates at either the optimal genotype (i.e., that with the maximum growth rate, circled in red) or a suboptimal peak (circled in cyan).

**Table 1.** *E. coli* data. Measured *IC*_50_ values (in *μg/ml*) and inferred growth rates (*r*) for *E. coli* exposed to 10^5^*μM* of Trimethoprim.

| *E. coli* | wild type | | GroEL+ | | $\Delta lon$ | |
| --- | --- | --- | --- | --- | --- | --- |
| Genotype | $IC_{50}$ | $r$ | $IC_{50}$ | $r$ | $IC_{50}$ | $r$ |
| Wt | 51.00 | 0.0212 | 46.85 | 0.0201 | 76.50 | 0.0273 |
| L28R | 1545.50 | 0.1740 | 788.50 | 0.1149 | 1000.00 | 0.1331 |
| A26T | 126.50 | 0.0372 | 29.50 | 0.0151 | 47.80 | 0.0204 |
| A26T,L28R | 1440.50 | 0.1667 | 861.00 | 0.1213 | 1866.50 | 0.1955 |
| P21L | 260.00 | 0.0580 | 206.50 | 0.0503 | 328.50 | 0.0670 |
| P21L,L28R | 1541.00 | 0.1737 | 644.50 | 0.1015 | 1348.50 | 0.1600 |
| P21L,A26T | 713.00 | 0.1080 | 952.50 | 0.1291 | 742.50 | 0.1107 |
| P21L,A26T,L28R | 1021.50 | 0.1348 | 1327.62 | 0.1585 | 2176.50 | 0.2150 |

**Table 2.** *L. grayi* data. Measured *IC*_50_ values (in *μg/ml*) and inferred growth rates (*r*) for *L. grayi* exposed to 10^5^*μM* of Trimethoprim. NA means the data were not available.

| *L. grayi* | wild type | | GroEL+ | | $\Delta lon$ | |
| --- | --- | --- | --- | --- | --- | --- |
| Genotype | $IC_{50}$ | $r$ | $IC_{50}$ | $r$ | $IC_{50}$ | $r$ |
| Wt | 10.35 | 0.0079 | 29.80 | 0.0152 | 45.50 | 0.0198 |
| L28R | 15.75 | 0.0103 | 149.00 | 0.0411 | 126.50 | 0.0372 |
| A26T | 5.59 | 0.0054 | 20.30 | 0.0120 | 36.50 | 0.0173 |
| A26T,L28R | 61.00 | 0.0237 | 188.50 | 0.0475 | 204.00 | 0.0499 |
| P21L | 41.84 | 0.0188 | 108.00 | 0.0337 | 96.35 | 0.0314 |
| P21L,L28R | 31.73 | 0.0158 | 38.10 | 0.0177 | 189.00 | 0.0476 |
| P21L,A26T | 13.62 | 0.0094 | 71.90 | 0.0262 | NA | NA |
| P21L,A26T,L28R | 10.50 | 0.0080 | 111.50 | 0.0344 | 29.00 | 0.0150 |

### Simulation Model

We simulated evolution on the 9 empirical fitness landscapes described above using DARPS (**D**iscrete **A**sexually **R**eproducing **P**opulation **S**imulator). DARPS was specifically designed to flexibly and efficiently simulate asexual reproduction and evolution of large populations of microbes on complex landscapes. During each discrete timestep, the number of individuals of each genotype grows exponentially according to its growth rate with stochastic single locus mutation, and then the entire population is reduced to the carrying capacity by frequency proportionate selection. We note that the classic Wright-Fisher model [25, 26] is a constant population size abstraction of the process implemented directly in DARPS. DARPS is described in more detail in [13] and open source code is available at [27].

**Table 3.** *C. muridarum* data. Measured *IC*_50_ values (in *μg/ml*) and inferred growth rates (*r*) for *C. muridarum* exposed to 10^5^*μM* of Trimethoprim.

| *C. miridarum* | wild type | | GroEL+ | | $\Delta lon$ | |
| --- | --- | --- | --- | --- | --- | --- |
| Genotype | $IC_{50}$ | $r$ | $IC_{50}$ | $r$ | $IC_{50}$ | $r$ |
| Wt | 3.15 | 0.0038 | 5.29 | 0.0052 | 9.28 | 0.0074 |
| L28R | 43.69 | 0.0193 | 118.50 | 0.0357 | 127.00 | 0.0373 |
| A26T | 4.43 | 0.0047 | 13.29 | 0.0093 | 13.60 | 0.0094 |
| A26T,L28R | 29.45 | 0.0151 | 174.00 | 0.0452 | 116.50 | 0.0353 |
| P21L | 2.15 | 0.0030 | 4.83 | 0.0050 | 4.40 | 0.0047 |
| P21L,L28R | 8.30 | 0.0069 | 21.25 | 0.0124 | 33.20 | 0.0163 |
| P21L,A26T | 2.43 | 0.0032 | 8.50 | 0.0070 | 11.65 | 0.0085 |
| P21L,A26T,L28R | 9.85 | 0.0077 | 19.75 | 0.0118 | 143.50 | 0.0402 |

For the simulation results reported here, mutation rates were assumed to be approximately 1 × 10^-10^ per locus per replication. In our DARPS model, the probability of mutation *P_m_* refers to one mutation in any of the 3 loci being studied per replication, so we used *P_m_* = 3 × 10^-10^. Bacterial population carrying capacity was set to *K* = 10^10^. Each simulation was initialized with a wild type population at carrying capacity, and allowed to run for up to 10^5^ timesteps, or for 1000 timesteps after *T_d_* (the timestep in which the genotype with the fastest growth rate became dominant), whichever came first. We then analyzed traces of the evolutionary dynamics to determine (i) which terminal genotype the population converged on, (ii) what evolutionary trajectory was followed from the wild type to the terminal genotype, (iii) when the terminal genotype dominated over 50% of the population (*T_d_*), and (iv) when the terminal genotype became “fixed” (*T_f_*), which we defined to be when it exceeded 99% of the population (due to ongoing mutational events, the terminal genotype will never comprise 100% of the population). We ran 1000 stochastic replicates of each simulation on the nine fitness landscapes.

### Within-path competition

We quantified the amount of within-path competition (*C_w_*) along the evolutionary trajectory followed in each of the simulations using the equation derived in [13], as follows:

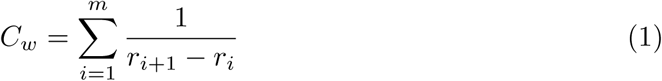

where *r_i_* represents the growth rate of genotype *i* along a trajectory comprising *m* steps from the genotype 1 (the wild type) to genotype *m* + 1 (the terminal genotype).

## Results

The topographies of all nine unique fitness landscapes are illustrated in terms of relative growth rates in Fig. 1, where the global peaks (fastest bacterial growth rates) are outlined in red and suboptimal peaks are outlined in cyan. Simulations of evolution on these DHFR landscapes demonstrate large differences in both the direction and speed of adaptive evolution, depending on the larger genotypic context. Although prior work on DHFR landscapes for the malaria parasite revealed that the “greediest” paths are not always those preferred by evolution [13], in these simulations we found that the greediest paths (shown by the thick blue trajectories in Fig. 1) were followed in all of the 1000 stochastic evolutionary simulations on each of these nine small landscapes. One representative simulation for each landscape is shown in Fig. 2. Below, we point out several notable findings in these results.

**Fig 2.**
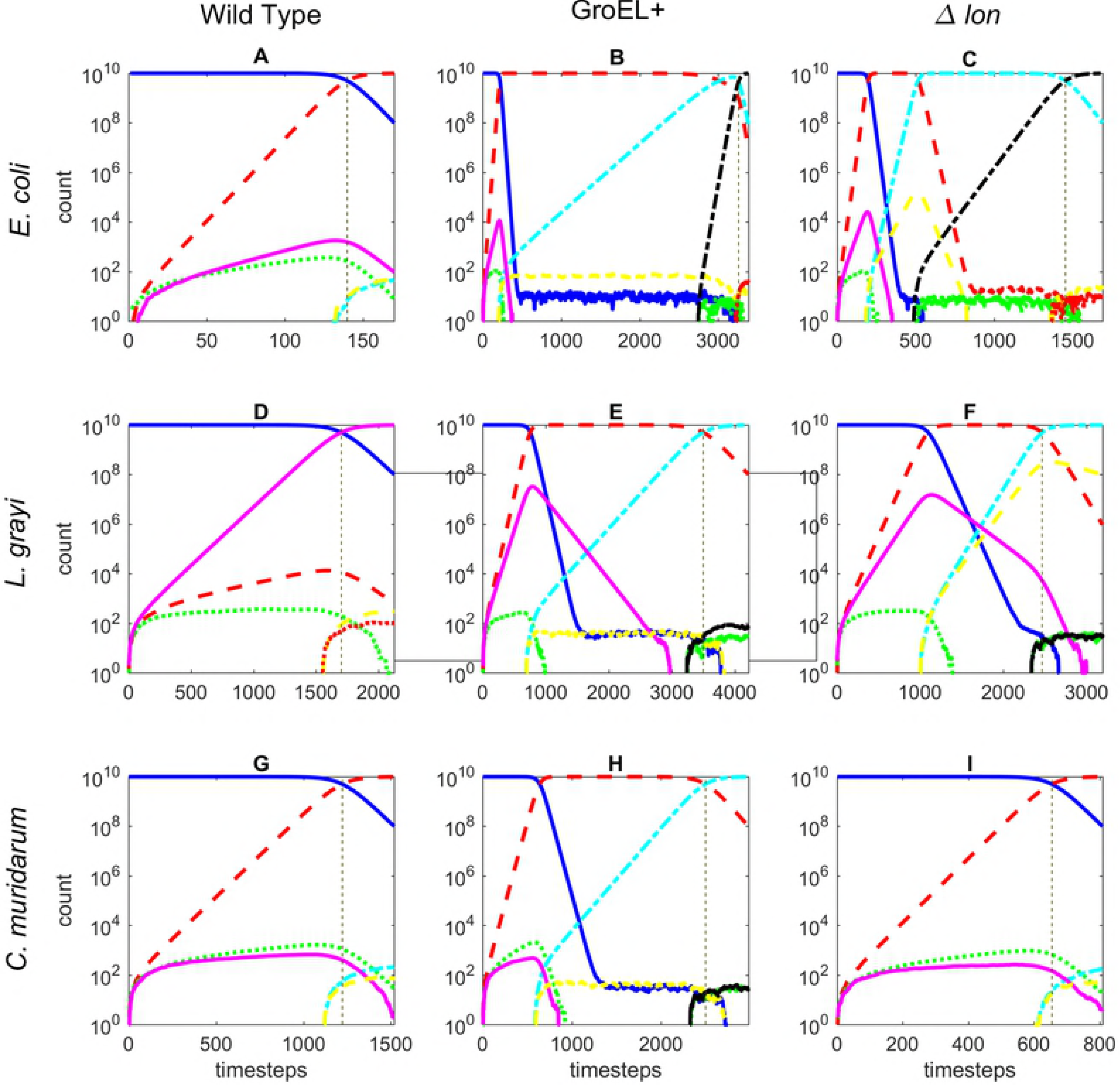
Simulation results. Representative evolutionary simulations on the nine empirical fitness landscapes for the three species (rows) with three genetic backgrounds (columns), starting from the wild type (WT). The vertical grey dashed line indicates the timestep at which the terminal genotype exceeded 50% of the population (*T_d_*) and the rightmost timestep shown is when it exceeded 99% of the population (*T_f_*). The legend in panel A applies to all panels. Note that the x-axes are scaled differently for each panel, and that y-axes are logarithmically scaled so that low-frequency genotypes are visible.

### Simulations of evolution demonstrate large differences in the direction of adaptive evolution across the nine fitness landscapes

#### Effect of PQC genotypic context on direction in *E. coli* landscapes

Within the *E. coli* landscapes (Figs. 1–2, top row, left to right), we observe that landscape topography changes across PQC backgrounds. In the wild type (WT) and the GroEL+ backgrounds, the population rapidly becomes fixed at the L28R optimal peak, which is only one mutational step from the initial (WT) genotype. In contrast, in the Δ*lon* PQC deletion background, the location of the optimal peak has shifted to P21L:A26T:L28R and evolution follows the WT→L28R→A26T:L28R→P21L:A26T:L28R pathway.

#### Effect of PQC genotypic context on direction in *L. grayi* landscapes

An intriguing pattern emerges in the *L. grayi* landscapes (Figs. 1–2, middle row, left to right). While all three PQC environments (WT, GroEL+ and Δ*lon*) have the same optimal peak at P21L:A26T:L28R, only the GroEL+ and Δ*lon* deletion landscapes reach this optimum, following the WT→L28R→A26T:L28R→P21L:A26T:L28R pathway. In the *L. grayi*/WT-PQC landscape, evolution instead takes the greedier step to the suboptimal peak of P21L, which becomes fixed. That is, despite the fact that all of the evolution is occurring on the same DHFR species backbone (*L. grayi*), the presence of the WT-PQC genomic background changes the topography, conferring a much different evolutionary outcome than the GroEL+ or Δ*lon* genomic backgrounds. Also note, that the two *L. grayi*/GroEL+-PQC and *L. grayi/*Δ*lon*-PQC landscapes have different suboptimal peaks (P21L for GroEL+; P21:L28R for Δ*lon*), which may not affect the preferred direction of evolution in these simulations, but might lead to different evolutionary outcomes under different conditions (environment, population genetic settings, etc.).

#### Effect of PQC genotypic context on direction in *C. muridarum* landscapes

In *C. muridarum* landscapes (Figs. 1–2, bottom row, left to right), the topography changes rather dramatically, with all three PQC landscapes having different optimal peaks. For the *C. muridarum*/WT-PQC landscapes, evolution proceeds along the single-step path to the optimal peak at L28R genotype. In the *C. muridarum/*Δ*lon* landscape the population also becomes fixed on the L28R genotype, but in this case this is a suboptimal peak that prevents evolution from reaching the optimal peak at P21L:A26T:L28R. In contrast, in *C. muridarum*/GroEL+ the population follows a two-step path to the optimal peak of A26T:L28R.

#### Effect of species-background on the direction of evolution

Just as the PQC genotypic context impacts the direction of evolution within each species (rows of Figs. 1–2), so does the species genotypic context impact the direction of evolution for each PQC profile (columns of Figs. 1–2). Note how the locations of the global peaks, the existence and location of suboptimal peaks, and the trajectories followed change across the landscapes as you look top to bottom within each column of Fig. 1.

#### Simulations of evolution demonstrate differences in the speed of adaptive evolution across the 9 fitness landscapes

We illustrate the average number of simulated timesteps it took for the terminal genotype to become dominant in the population (*T_d_*) in Fig. 3, due to the topographic differences if the landscapes shown in Fig. 1. Note that evolution to the terminal genotype is universally slower in the *L. grayi* DHFR background than the other two species backgrounds, but that the relative evolutionary speeds in *E. coli* and *C. muridarum* depend on the PQC genotypic context.

It is also important to note how the overall growth rates of the genotypes in a landscape do not govern the speed of evolution across a landscape. For example, although the *E. coli* growth rates are much higher than those of *L. grayi* and *C. muridarum* across all PQC backgrounds (Fig. 1 and Table 1), evolution does not always proceed fastest along these landscapes (Fig. 3). This is an important reminder that the speed of evolution is not a function of the fitness of individual genotypes, but is largely governed by the differences in fitnesses of adjacent genotypes in an evolutionary trajectory [13], as quantified by the within-path competition (*C_w_*) shown in Eq. (1). For example, in these simulations *T_d_* is shown be a slightly sublinear function of Cw (Fig. 4, *R*^2^ > 0.99, *p* ≈ 0). Despite the stochastic nature of these simulations, *T_d_* can be seen to be very consistent across the 1000 repetitions of each simulation of these very large populations (Fig. 4). The competition values along each individual step of the trajectories followed are shown in blue in Fig. 1, with the total *C_w_* for each trajectory shown above each landscape in Fig. 1.

**Fig 3.**
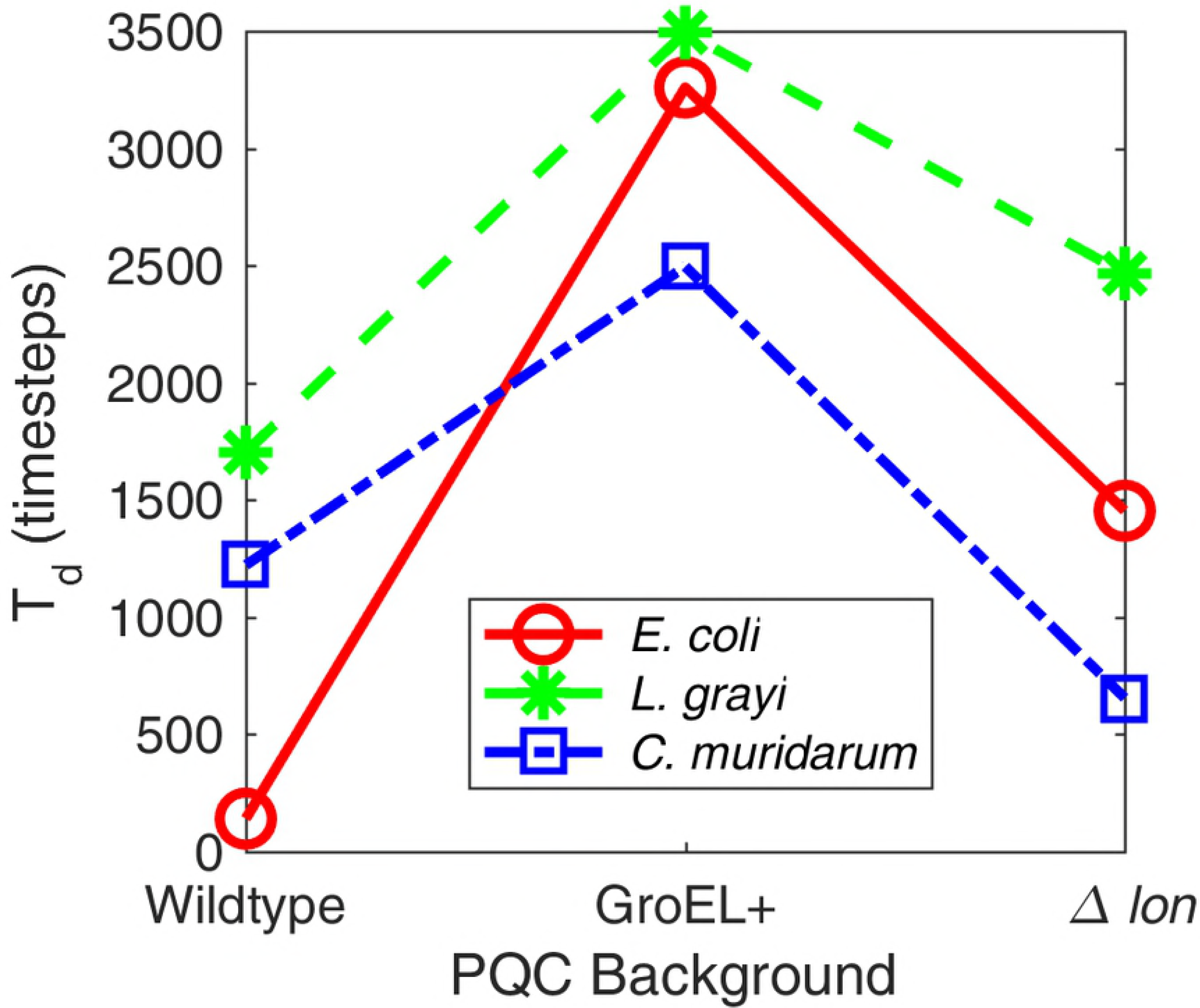
Speed of evolution. The number of timesteps required for the terminal genotype to exceed 50% of the population (*T_d_*) varies greatly for different PQC genotypics contexts and different species.

**Fig 4.**
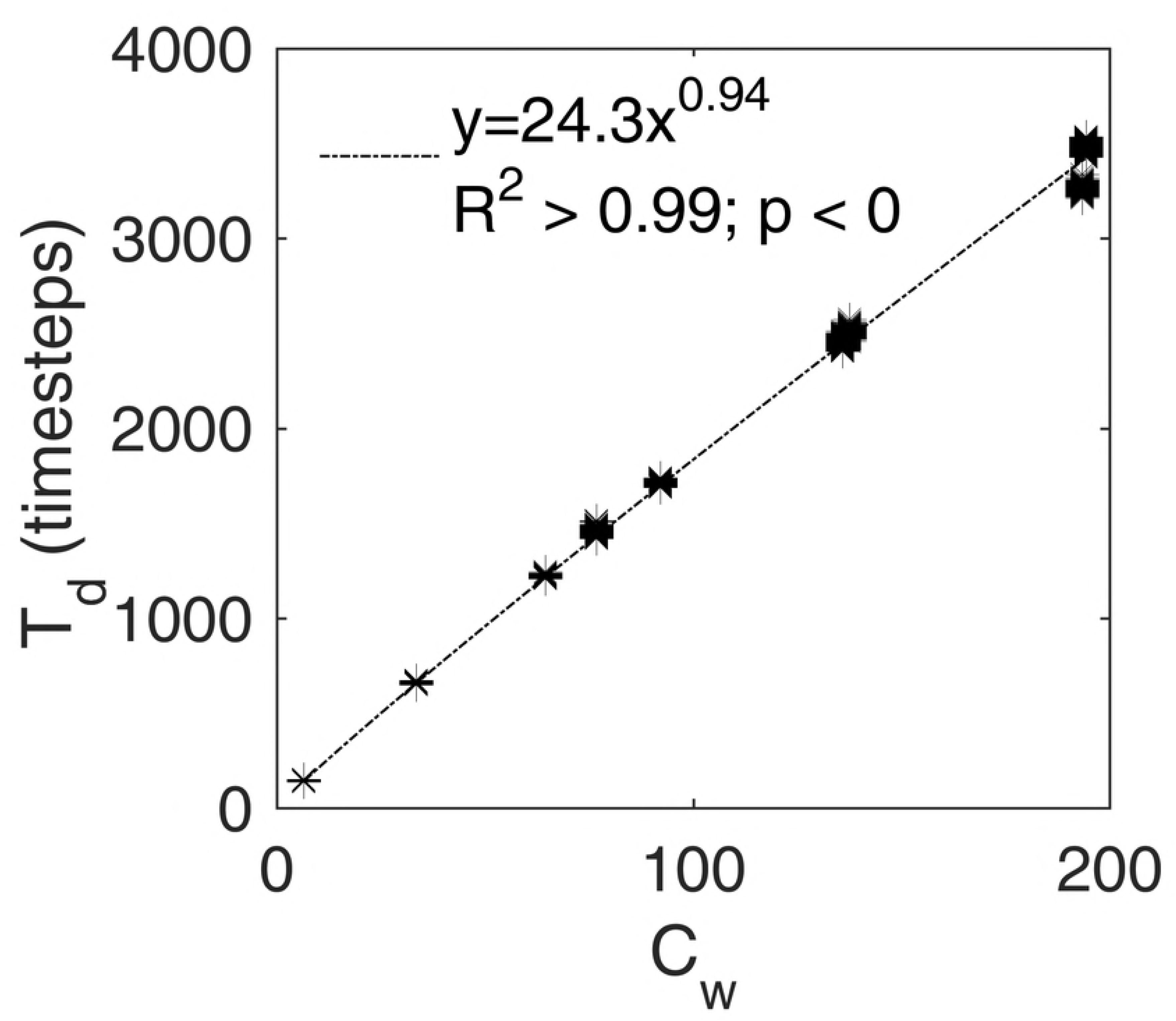
Within-Path competition governs evolutionary speed. The amount of within-path competition (*C_w_*) governs the number of timesteps required for the terminal genotype to exceed 50% of the population (*T_d_*).The asterisks represent observations from 1000 simulations on each of the 9 fitness landscapes, and the best-fit curve was determined by regressing *log*(*T_d_*) *vs. log*(*C_w_*).

## Discussion

### Genotypic context alters fitness landscape topography

In this study, we have identified that genotypic context alters fitness landscape topography for antibiotic resistance, which in turn influences 3 aspects of evolutionary dynamics: (i) the distribution of optimal and suboptimal peaks on a fitness landscape, (ii) the “preferred” direction of adaptive evolution and (iii) the speed at which said evolution occurs.

### Species differences in protein backbone alters the speed and direction of evolution

Evolutionary simulations on these landscapes illustrate how evolution occurs differently across species. With regard to the evolution of drug resistance, these findings indicate that even subtle differences in the amino acid sequence for otherwise conserved enzymes can have a powerful effect on how evolution occurs (both speed and direction). This implies that we cannot assume that even closely related microbial pathogens will evolve resistance to drugs using the same evolutionary trajectory, as the fitness landscape underlying resistance may be different. This might be complicating news for the burgeoning field of resistance management: instead of being able to adopt a one-size-fits-all approach to managing resistance, we may have to engineer our managements to very specific genotypic contexts.

### Differences in protein quality control alter the speed and direction of evolution

Two thirds of the landscapes with PQC modifications had longer evolutionary trajectories than in the wild type, and in *L. grayi* they even had different initial directions. This supports the idea that global protein quality control regulation may have prescribed ways of altering the landscape, maybe related to the way they influence the lifetime and performance of enzymes in a cell [21, 28]. In this setting, mutations may alter resistance patterns not because they affect the way a drug binds but because they affect the interaction between a protein effector and the PQC machinery. This would suggest a mechanism for how resistance in microbes can be so biochemically and biophysically diverse, even in well-characterized systems like DHFR and antifolates: an enzyme might avoid the effects of a drug through altering its interaction with other genes maybe even in lieu of altering the binding of a antibiotic drug.

### General note on the speed of evolution

The speed of evolution from the wild type genotype to the terminal genotype in evolutionary simulations is shown to vary greatly across different genotypic contexts, and in a manner that is not related to the absolute fitnesses of the nodes in the respective landscapes. More broadly, this study affirms the relationship between within-path competition (*C_w_*, defined by Eq. (1)) and the speed of evolution (determined via simulation) [13]. As a general observation, studies that examine the speed of evolution have been all but ignored in the study of empirical fitness landscapes, although it has recently been demonstrated to be an important property of evolutionary dynamics [13]. In particular, discussions that invoke empirical fitness landscapes in discussing how one might better prevent or manage drug resistance in plant and animal infectious disease should be especially mindful of the speed of evolution: True resistance management should not only consider which pathways evolution will traverse towards maximal resistance, but how fast certain pathways might occur relative to others.

Our findings highlight that, like the “preferred” direction of evolution, the speed of evolution should be considered in any study that examines how and why fitness landscape topography determines evolutionary outcomes.

### Conclusions

In closing, we have revealed how an under-appreciated determinant of the topography of an adaptive landscape – the larger genotypic context outside the specific target genes being studied – influences the speed and direction of adaptive evolution using empirical data and computer simulations. The findings of this study have broad implications for public health, technology, and theory regarding speciation and evolvability. In the context of public health and biomedicine, our results imply that even subtle genetic differences between microbial populations can be sufficient to drive different evolutionary outcomes, both in terms of the predicted speed and direction of evolution. This suggests that future efforts at “resistance management” need to consider very specific genomic and genetic details about the population being managed before rigorous and effective management strategies are engineered.

In addition, our results highlight how particular “off target” mutations (in our study, PQC modifications) can have powerful influences on evolutionary outcomes. Consequently, genomic screens for “resistance mutations” should focus on potential signals across the genome, rather than a singular focus on genes that are the presumptive target of therapy. Our results illustrate that there are multiple ways to subvert the effects of a drug, sometimes involving genes and gene networks that are not intuitively (or biophysically) linked to the phenotype of interest (in this case, protein quality control genes having no specific connection to DHFR activity). Similarly, our results underscore the potential perils of engineering mutations associated with a given phenotype into different genomic backgrounds, as in CRISPR-mediated genetic engineering. In such scenarios, differences in genomic background of strains in which a given SNP is being engineered can not only influence the effect of the mutation being introduced, but also, the downstream evolution of different populations.

Lastly, our results speak to the notion that small genetic differences between populations may be sufficient to induce larger downstream divergence, eventually leading to speciation. Specifically, our study is consistent with the expectation that reproductive isolation arises rapidly in rugged fitness landscapes (e.g., in a Bateson-Dobzhansky-Muller framework [29], or holey landscape [30]). By examining genetic differences at various scales (single nucleotide polymorphisms in target resistance genes, species-specific differences in genetic background, and changes to off-target genes), we demonstrate how “difference can beget difference” in Darwinian evolution, affecting both the degree and rate of divergence.

## Acknowledgments

The authors would like to thank S. Scarpino and J. Rodrigues for helpful discussions on the project. C.B. Ogbunugafor would like to acknowledge NSF RII Track-2 FEC (Award Number: 1736253), “Using Biophysical Protein Models to Map Genetic Variation to Phenotypes” for funding support.

